# Myelin water imaging depends on white matter fiber orientation in the human brain

**DOI:** 10.1101/2020.03.11.987925

**Authors:** Christoph Birkl, Jonathan Doucette, Michael Fan, Enedino Hernandez-Torres, Alexander Rauscher

## Abstract

The multiexponential T_2_ decay of the magnetic resonance imaging (MRI) signal from cerebral white matter can be separated into short components sensitive to myelin water and long components related to intra- and extracellular water. In this study we investigated to what degree the myelin water fraction (MWF) depends on the angle between white matter fibers and the main magnetic filed. Maps of MWF were acquired using multi echo CPMG and GRASE sequences. The CPMG sequence was acquired with a TR of 1073 ms, 1500 ms and 2000 ms. The fiber orientation was mapped with diffusion tensor imaging. By angle-wise pooling the voxels across the brain’s white matter, an orientation dependent MWF curve was generated. We found that MWF varied between 25% and 35% across different fiber orientations. The orientation dependency of the MWF is characterized by a dipole-dipole interaction model. Furthermore, the selection of the TR influences the orientation dependent and global white matter MWF. White matter fiber orientation induces a strong systematic bias on the estimation of MWF. This finding has important implications for future research and the interpretation of MWI results in previously published studies.

## Introduction

Myelin is a lipid bilayer membrane wrapped around the axon, protecting it from mechanical and chemical damage (1, 2) and facilitating fast saltatory signal conduction (3, 4). The space between the myelin bilayers is filled with water which is commonly referred to as myelin water. Due to reduced mobility within this confined space, myelin water has a magnetic resonance T_2_ T_2_ relaxation time of around 10 ms at a field strength of 3T. In contrast, the T_2_ relaxation time of the intra- and extracellular water is in the range of 30 ms to 60 ms, and the cerebrospinal fluid (CSF) has a T_2_ relaxation time of more than one second (5, 6). The resulting multiexponential T_2_ signal decay of a white matter voxel can be measured using a multi-echo spin echo scan and decomposed into its individual components, resulting in a T_2_ spectrum (7). The area under the short T_2_ components, referred to as the myelin water peak, divided by the total area is defined as the myelin water fraction (MWF). A strong correlation between the MWF and independent measures of myelin has been demonstrated in various post-mortem studies (8–10). Over the past two decades, myelin water imaging (MWI) has been applied to mild traumatic brain injury (11), aging (12), spinal cord injury (13), neonates (14), and multiple sclerosis (15–18), among others. The human brain’s white matter is highly anisotropic. This circumstance is widely exploited in diffusion tensor imaging (DTI), which allows to map the brain’s complex fiber architecture (19). The orientation of white matter fibers with respect to the main magnetic field B_0_ also affects the magnitude and phase of the complex MRI signal and a wide range of quantitative MRI parameters. Strong orientation effects have been reported for 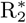 relaxation (20–25), gradient echo phase (21, 24, 26), quantitative susceptibility maps (QSM) (27). A weak orientation effect also exists for T_1_ relaxation (28, 29). Moreover, due to anisotropy of the cerebral vascular architecture, dynamic susceptibility contrast (DSC) perfusion measurements conducted with both gradient echo (30) and spin echo sequences (31) show an orientation dependent behaviour. The orientation dependency of the gradient echo signal of white matter is extensively studied and is ascribed mainly to the anisotropy of the magnetic susceptibility caused by the myelinated nerve fibers (24, 25, 32, 33). The observation that the spin echo DSC signal (31) and white matter R_2_ (25) also depend on the fiber orientation suggests that MWI is potentially affected as well. Therefore, we investigated whether and to what degree the MWF depends on the angle between the white matter fiber tracts and the main magnetic field B_0_. In the present study, we acquired MWI using a gradient and spin echo (GRASE) sequence (34), which is widely used in literature, and the Carr-Purcell-Meiboom-Gill (CPMG) sequence, which is the gold standard. The CPMG sequence was further acquired at various repetition times (TR). We show that the MWF strongly depends on white matter fiber orientation, independent of MRI sequence.

## Methods

Eight healthy volunteers (3 female, 5 male) with a mean age of 26 years (age range = 21 − 33 years) and without any history of neurological disorder participated in this study, which was approved by the University of British Columbia Clinical Research Ethics Board. All volunteers gave written informed consent. MR imaging was performed on a 3T MR system (Ingenia Elition, Philips Medical Systems, Best, The Netherlands) using a 32-channel SENSE head coil.

### MRI acquisition

A 3D T_1_ weighted sequence with echo time (TE) = 4.9 ms, TR = 9.8 ms, flip angle = 8°, a resolution of 0.8 × 0.8 × 0.8 mm^3^, a compressed SENSE (CS) factor of 3.6 and acquisition time of 5:50 min was acquired for anatomical overview. CS is a combination of compressed sensing (35, 36) and SENSE (37). For the calculation of fiber orientation a DTI sequence with TE = 60 ms, TR = 4111 ms, b-value = 700 s/mm^2^, 60 diffusion directions, a resolution of 2.3 × 2.3 × 2.4 mm^3^, multi band factor 2 (38), SENSE factor 2.1 and acquisition time of 4:19 min was acquired. MWI was performed using following sequences: (I) a GRASE sequence with TR = 1073 ms, SENSE of 2.5, three gradient echos per spin echo and acquisition time of 11:29 min. (II) a CPMG sequences with TR = 1073 ms and acquisition time of 09:43 min. (III) a CPMG sequence with TR = 1500 ms and acquisition time of 13:33 min. (IV) a CPMG sequence with TR = 2000 ms and acquisition time of 18:04 min. All CPMG sequences were accelerated with a CS factor of 7 and all MWI sequences had 48 echoes with the first echo at TE = 8 ms, ΔTE = 8 ms, and a spatial resolution of 0.96 × 0.96 × 2.5 mm^3^. 3D T_1_, DTI and MWI sequences were acquired in axial orientation without angulation. Due to the long data acquisition times, not all sequences were acquired in all volunteers.

### Image analysis

DTI data were analysed with the FMRIB Software Library (FSL v5.0.9)(39–41). Distortions induced by eddy currents and head motion were corrected by FSL’s eddy_correct. FSL DTIFIT was used to calculate the diffusion tensor model and thus the eigenvalues and eigenvectors. For MWI analysis, the T_2_ distributions were computed using a regularized non-negative least squares (NNLS) algorithm with stimulated echo correction and a T_2_ range of 8 ms to 2.0 s (6, 34, 42, 43). The multi-exponential signal decay is expressed as a T_2_ distribution where the myelin water component is defined as the T_2_ times of the distribution between 8 ms and 25 ms. The intra- and extra-cellular water component is defined as the T_2_ times above 25 ms. The cut-off between the two water pools was set to 25 ms, which was based on the measured T_2_ distributions. The MWF was calculated as the ratio of the myelin water T_2_ components to all T_2_ components. The T_1_ and DTI data was linearly registered to the MWI using FLIRT. FSL’s vecreg was used to register the first eigenvector to the MWI space. The fiber orientation was calculated from the angle between the first eigenvector to the direction of the main magnetic field B_0_ (21). Brain extraction was performed using FSL’s brain extraction tool (bet) and FSL’s FAST was used to generate a white matter mask, which was eroded with a 3 × 3 × 3 kernel. The MWF, myelin water T_2_ and intra- and extra-cellular water T_2_ were computed as a function of local fiber orientation *θ*. The fiber orientations were devided into 18 intervals of 5° between 0 ° (parallel to B_0_) and 90° (perpendicular to B_0_). To compute MWF(*θ*), voxels from across the entire white matter were pooled for each angle interval, in order to suppress the influence of tract specific differences in MWF.

### Fit of the dipole interaction model

Highly ordered biological tissues give rise to orientation dependent T_2_ signal via dipole-dipole interaction (44). Therefore, a dipole-dipole interaction model (25) according to

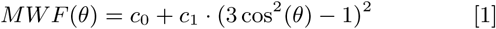

was fitted to the MWF data. c_0_ represents the orientation independent part of the MWF, c_1_ a parameter scaling the orientation dependent part of the MWF and *θ* is the angle between white matter fibers and the main magnetic field B_0_. Fitting was performed using MATLAB’s nonlinear least squares algorithm.

### Statistical analysis

All parameters were tested for normal distribution using the Shapiro-Wilk test. A t-test was used to test significance of differences between GRASE and CPMG. To test whether the fibre angle and TR had a significant effect on MWF and T_2_, an ANOVA was used. All statistical analyses were performed using R (version 3.5.1, The R Foundation for Statistical Computing).

## Results

Representative maps of the angle between white matter fibers and B_0_, and the corresponding MWF maps are shown in Figure 1. Overall, the GRASE and CPMG sequence provide comparable results, although using a CPMG sequence resulted in higher MWF.

**Fig. 1.**
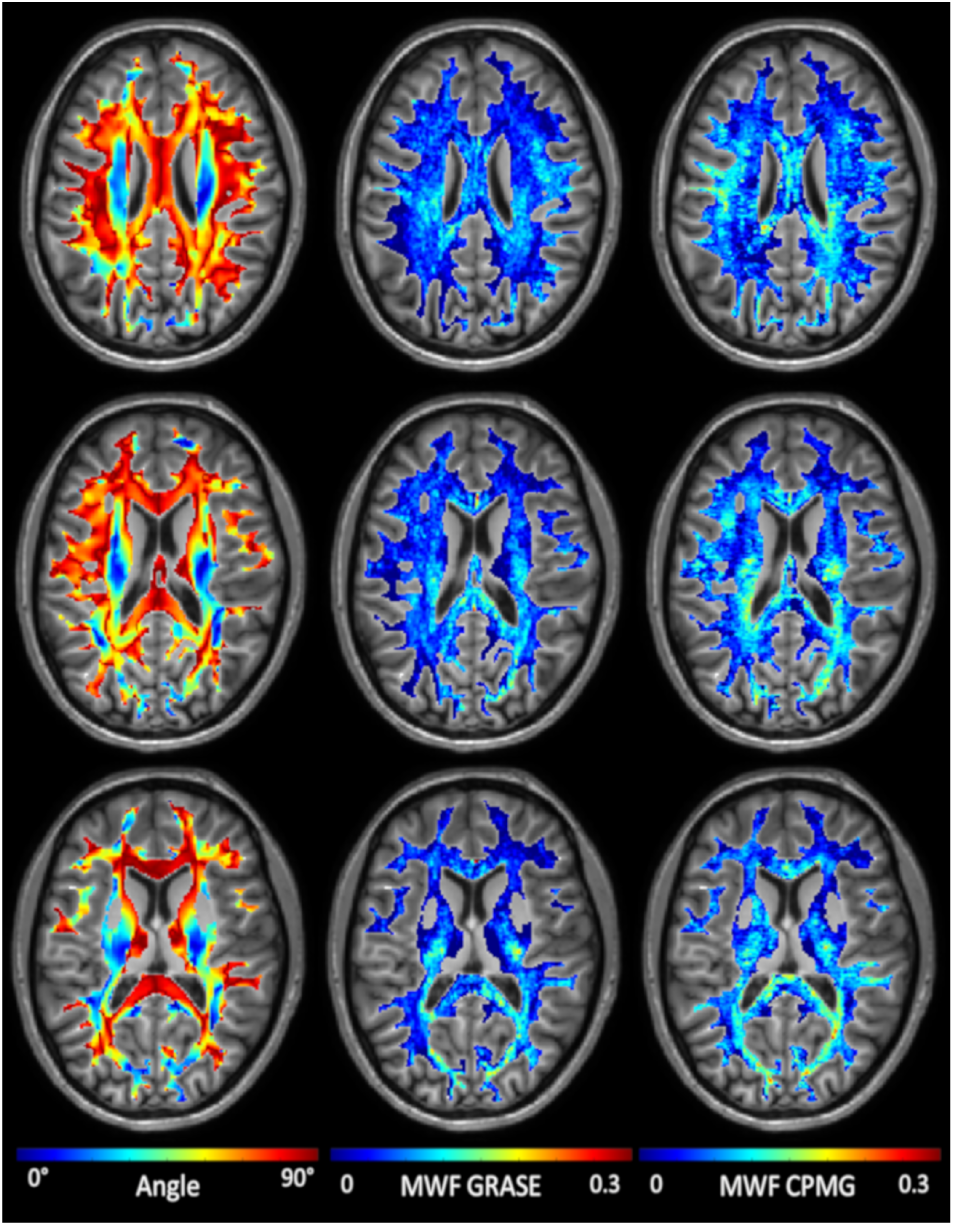
Representative fiber angle and MWF maps, acquired using a CPMG and GRASE sequence, of a single subject.

The relationship between MWF, myelin water T_2_ and intra- and extra-cellular water T_2_ and the white matter fiber orientation, is shown in Figure 2. In general, the MWF decreased with increasing fiber angles and reached a minimum between 50°and 60°, followed by an increase towards angles of 90°. The solid red line represents the fit of the dipole-dipole interaction model to the measured MWF data (Figure 2A). The MWF varied by approximately 35% for the GRASE sequence, and by approximately 22% for the CPMG sequence, across different fiber orientations. Using a GRASE sequence resulted in a stronger MWF orientation dependency, as evident by a lager c1 coefficient (see Table 1). A summary of all fit parameters is given in Table 1. Myelin water T_2_ (Figure 2B) and intra- and extra-cellular water T_2_ (Figure 2C) increased with increasing fiber angle up to a maximum between 15°to 25° followed by a decrease. Between GRASE and CPMG, there was a significant difference in the orientation dependency of the MWF (p = 0.002), the myelin water T_2_ (p < 0.001) and no significant difference in the orientation dependency of the intra- and extra-cellular water T_2_ (p = 0.79).

**Table 1.** Summary of the dipole-dipole fits to the measured orientation dependent MWF. c_0_ represents the orientation independent part of the MWF. c_1_ is a constant scaling the orientation dependent part of the MWF.

| Sequence | $TR$ (ms) | $c_0$ | 95% CI | $c_1$ | 95% CI | SE |
| --- | --- | --- | --- | --- | --- | --- |
| GRASE | 1073 | 0.062 | (0.060-0.065) | 0.010 | (0.008-0.011) | 0.0049 |
| CPMG | 1073 | 0.060 | (0.059-0.062) | 0.005 | (0.005-0.006) | 0.0012 |
| CPMG | 1500 | 0.065 | (0.063-0.067) | 0.004 | (0.003-0.005) | 0.0018 |
| CPMG | 2000 | 0.058 | (0.056-0.060) | 0.003 | (0.001-0.004) | 0.0015 |

**Fig. 2.**
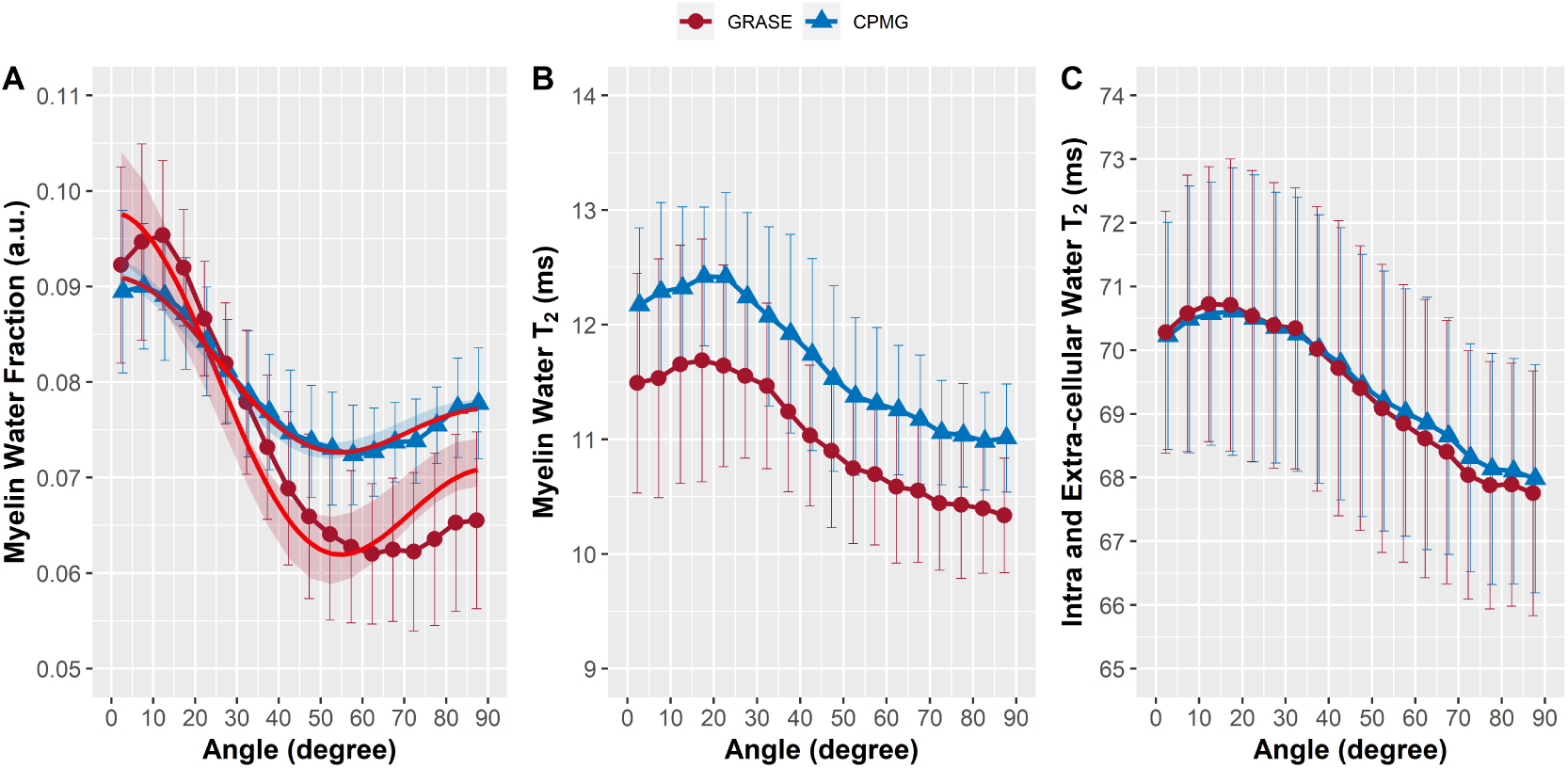
Myelin water fraction (MWF) (A), myelin water T_2_ (B) and intra- and extra-cellular water T_2_ (C) as function of fiber orientation acquired using a GRASE (red) and CPMG (blue) sequence, both with TR = 1073 ms. The solid red lines represent the fits of the dipole-dipole interaction model to the MWF data. The minimum of the orientation dependent MWF was between 50°and 60°, which agrees well with the minimum of the dipole-dipole model at 54.7°. The shaded area represents the 95% confidence interval.

### Effect of TR

With increasing TR from 1073 ms to 1500 ms and 2000 ms, the orientation dependent MWF decreased across all regions, as shown in Figure 3. In general the TR of the CPMG sequence had a significant influence on MWF (p < 0.001), myelin water T_2_ (p < 0.001) and intra- and extra-cellular water T_2_ (p < 0.001) as shown in Figure 4. The shape of the orientation dependent MWF, myelin water T_2_ and intra- and extra-cellular water T_2_ was similar but shifted with increasing TR. The effect of TR is further evident by plotting the global white matter mean of the MWF (Figure 5A), myelin water T_2_ (Figure 5B) and intra- and extra-cellular water T_2_ (Figure 5C). For example the MWF decreased by 21.5% from 0.079 at TR = 1073 ms to 0.062 at TR = 2000 ms (p = 0.006) as shown in Figure 5A.

**Fig. 3.**
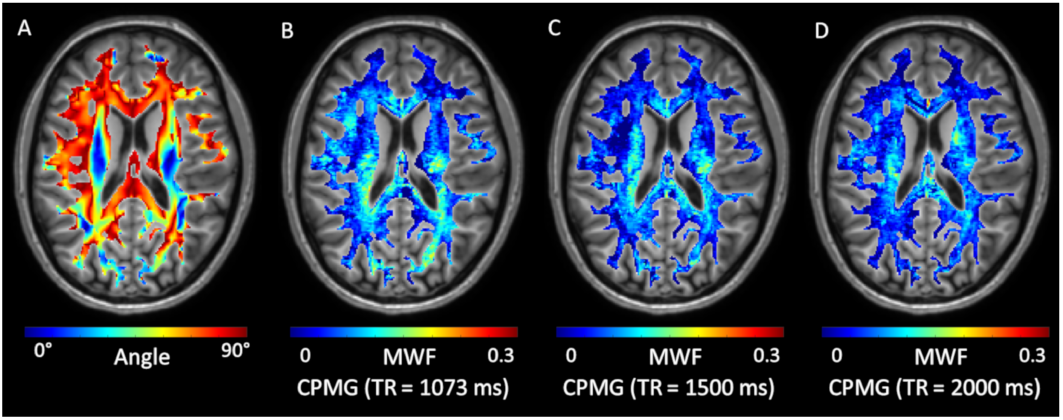
Representative fiber angle and MWF maps acquired using a CPMG sequence with TR = 1073 ms, 1500 ms and 2000 ms. Overall, MWF decreases with increasing TR.

**Fig. 4.**
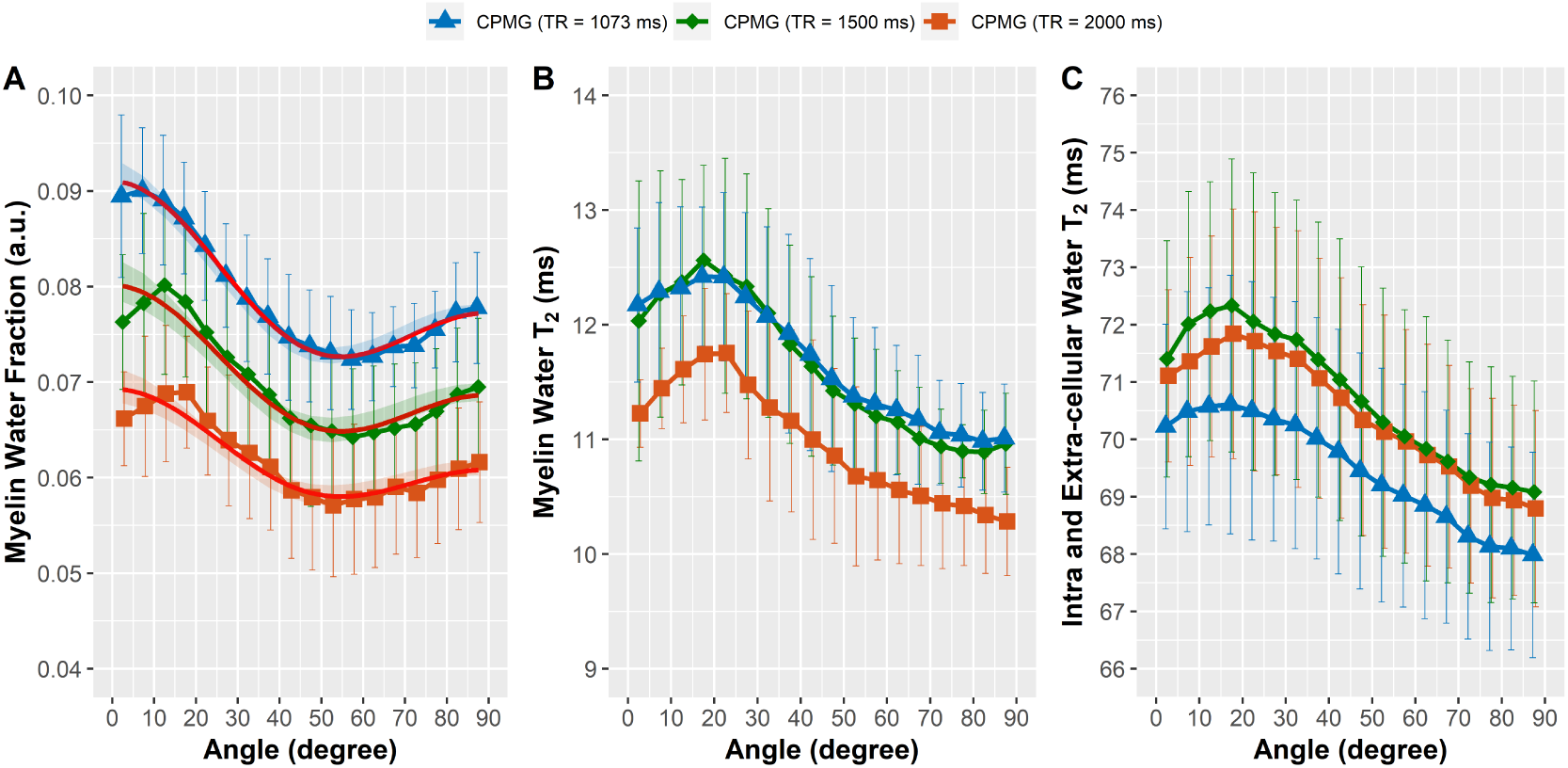
Myelin water fraction (MWF) (A), myelin water T_2_ (B) and intra- and extra-cellular water T_2_ (C) as function of fiber orientation acquired using a CPMG sequence with a TR of 1073 ms (blue), 1500 ms (green) and 2000 ms (orange). The solid red line represents the fit of the dipole-dipole interaction model to MWF data. The minimum of the orientation dependent MWF was between 50°and 60°, which agrees well with the minimum of the dipole-dipole model at 54.7°. The shaded areas represent the 95% confidence intervals.

**Fig. 5.**
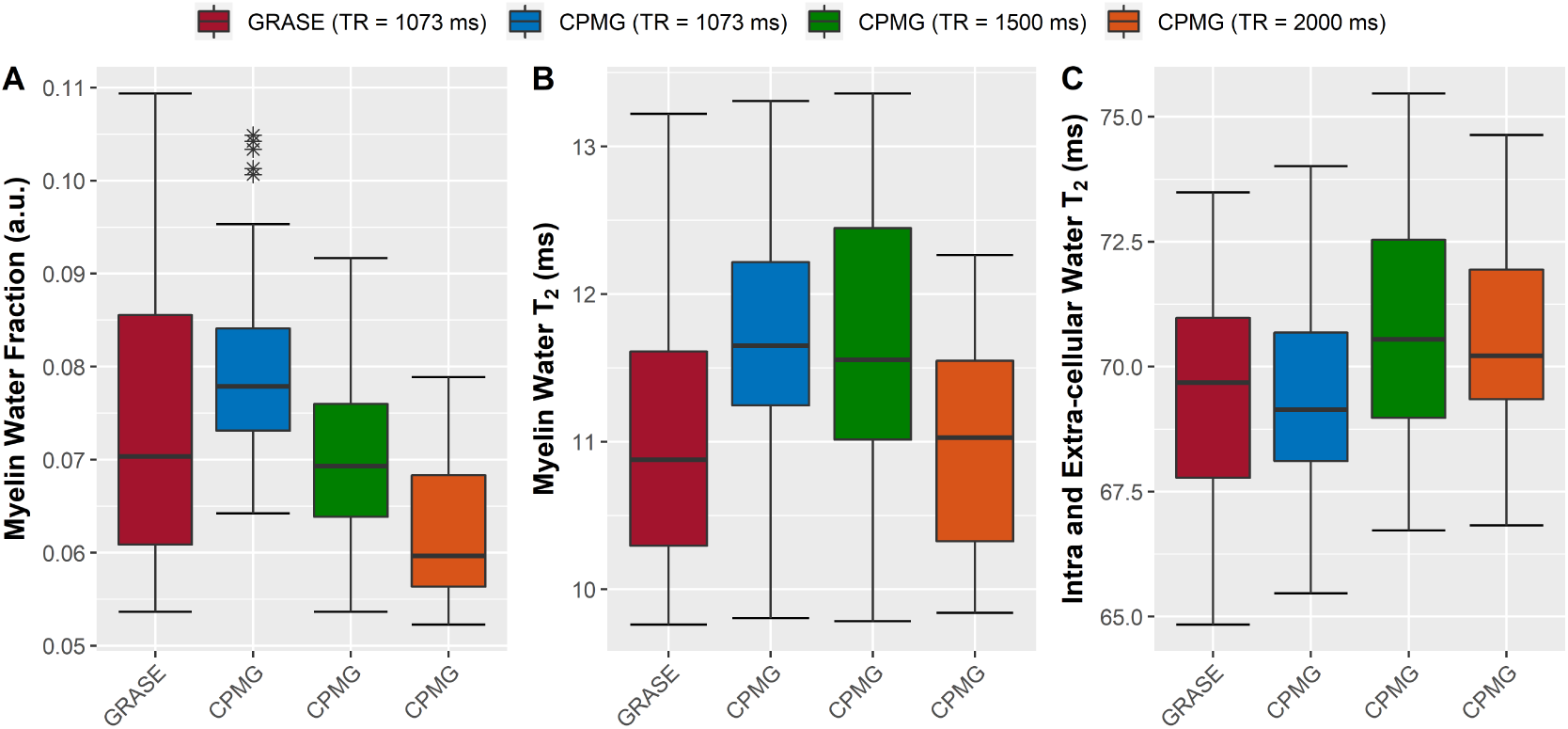
Global white matter MWF (A), myelin water T_2_ (B) and intra- and extra-cellular water T_2_ (C). TR has a significant impact on MWF (p < 0.001), myelin water T_2_ (p < 0.001) and intra- and extra-cellular water T_2_ (p < 0.001).

## Discussion

We demonstrated that the measurement of MWF is considerably influenced by the angle between white matter fiber tracts and the main magnetic field. Furthermore, we showed that the TR of the sequence has an impact on estimating the MWF. Orientation effects of the transverse relaxation have been mainly ascribed to dipole-dipole interaction and to magnetic susceptibility effects. Orientation dependency of T_2_ has been described in vivo in highly ordered tissues, such as cartilage and tendons (45, 46). Such structures give high T_2_ signal when they are oriented at angles where the term 3 cos^2^(*θ*) − 1 describing the z-component of the dipole field is zero. Another cause of orientation effects is the magnetic susceptibility of tissue, which has a strong influence on phase and magnitude of the gradient echo signal. While the orientation dependency of white matter 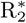 has been studied extensively (20–22, 24, 25, 47), the effect of tissue orientation on R_2_ in the brain has been considered weak, with the notable exception of spin echo DSC MRI, where a paramagnetic contrast agent amplifies the magnetic field inhomogeneities around blood vessels. Diffusion in the vicinity of the anisotropic white matter vasculature gives rise to a strong orientation dependency in the measurements of cerebral blood flow and volume (31). Of course, such orientation effect is also present and much stronger in gradient echo DSC (30), where the signal is dominated by static dephasing. Oh and colleagues reported a weak orientation dependency of R_2_ in ex vivo formalin fixed white matter that is best described by dipole-dipole interaction (25). Based on their finding, the authors suggested that MWF should also weakly depend on fiber orientation. However, the angle effects in their R_2_ data were small, which could be a consequence of fixation induced cross-linking, which reduces tissue anisotropy at length scales that are relevant to T_2_ relaxation. Furthermore, with the decreased T_2_ relaxation times due to tissue fixation (48) and due to the high field strength of 7T, the shortest echo time of 9 ms may be too long to capture orientation effects under these conditions. In the present work in vivo at 3T and with 8 ms echo spacing, we saw that MWF was approximately 30% lower at angles between 50° and 60° compared to tissue parallel to B_0_. All MWF measurements followed this pattern, with a minimum near the magic angle of 54.7°. Fitting the dipole-dipole model to our data resulted in good fit quality, in particular for the CPMG data. Note that there is almost perfect overlap between the average CPMG MWF(*θ*) and the dipole-dipole model which is reflected in the low standard error of the regression. The orientation behaviours of GRASE and CPMG were different, in particular at larger angles. The attenuated recovery of the MWF in GRASE at angles above 54.7°compared to the more pronounced recovery in the CPMG may be due to some 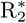 weighting of the GRASE sequence, which may lead to a reduction of the signal due to field inhomogeneities created by the myelinated axons at larger angles. The dependency of MWF on TR could be explained by the different T_1_ relaxation times of myelin water and the intra- and extracellular water. The overall T_1_ relaxation of white matter at 3T is around 1100 ms (49, 50), whereas T_1_ of myelin water is around 200 ms (51–53). Therefore, myelin water is almost fully relaxed even at the shortest TR used in the present study, whereas the intra- and extracellular water signal is attenuated and increases with increasing TR. As a result, the MWF decreases with increasing TR. Schyboll and colleagues showed a small orientation dependency in both T_1_ and water content measurements, of 2.5% and 0.8%, respectively (28). Interestingly, the shape of the water content curve mirrors the MWF curve in our study. In a very recent paper, the same authors showed that the susceptibility effects of the myelin sheath only cause a very weak orientation dependency in T_1_ of about 0.4 × 10^−4^*Hz* between parallel fibers and fibers at the magic angle (29).

The good agreement between MWF(*θ*) and the dipole-dipole model and previous observations in other ordered tissues that are also described by dipole-dipole interaction suggest that this interaction is a main cause of orientation effects in MWF. The question is how the dipole-dipole interaction translates to the observed MWF(*θ*). A direct effect of dipole-dipole interaction in the myelin water pool itself would result in the reverse behaviour, with a maximum at the magic angle instead of the observed minimum. Henkelman et al. (46) argued that the alignment of water molecules with filaments but not with cylindrical or plane surfaces would cause dipole-dipole coupling effects in T_2_. Such effect is seen in cartilage, where the collagen fibers lead to a two to threefold increase in T_2_ signal at the magic angle (54). Filaments are also present within the axon as microtubule and neurofilaments (55), which may be able to produce a magic angle effect in the T_2_ signal of white matter. This interpretation is in agreement with the reduced orientation dependency of T_2_ in ex vivo formalin fixed tissue (25), where cross-linking reduces the anisotropy. Additional controlled experiments in tissue samples and animal models are needed to fully explain the origins of orientation dependency in MWF.

This study has limitations. We only investigated TR up to 2000 ms, which is approximately two times the T_1_ of the long T_1_ component. The shortest TR of 1073 ms, on the other hand, was defined by the specific absorption rate of the scan. With the required anatomical and DTI scans and the comparison with the GRASE approach at 1073 ms, the total scan protocol took around 75 minutes. Therefore, the present study is not able answer the question how MWF behaves in the absence of T_1_ weighting. A comprehensive study that explores the effects of TR would require several long scans with TRs of up to 5000 ms, where T_1_ weighting of the intra- and extra-cellular water is reduced to less than 2%. Such study may be feasible with reduced brain coverage and lower spatial resolution, which both make image registration more challenging, and by not scanning every TR in every participant.

While the model of dipole-dipole interaction fits the data well, additional effects may play a role in the observed MWF behaviour. The approach of using DTI to determine the fiber orientation and pooling voxels according to their local fiber orientation is widely used and accepted (20, 21, 24, 30, 56). Since for each angle interval voxels are pooled from across the entire brain, potential tract specific differences in MWF get averaged out. Moreover, MWF(*θ*) of all subjects followed this pattern, independently of individual differences in head orientation or white matter anatomy. Finally, if anatomical differences played a major role, MWF(*θ*) would exhibit an irregular pattern instead of following a smooth and simple trigonometric function with only one angle dependent parameter.

Our findings have several consequences for the interpretation of past results and for future research. The orientation dependency of MWF might mask longitudinal changes in MWF or changes between cohorts, if whole WM averages are investigated. A corresponding effect was observed in white matter R_2_*, where only an orientation dependent analysis was able to reveal group differences (47, 56). Kor et al. modelled the 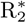 relaxation as a function of white matter orientation, myelin, and iron content and were able to compute whole white matter myelin and iron content (47). This approach assumed that the effects of iron are orientation independent and that only the myelin sheath gives rise to an orientation dependent 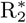. Since iron contributes to more than 25% of the apparent MWF (57), an orientation dependent analysis accompanied by a model that includes the effects of both myelin and iron may shed further light on the orientation dependency of MWF.

Our findings suggest that, when comparing MWF of individual subjects to a myelin water atlas (58, 59), these atlases have to be updated with DTI data. In particular, fiber tracts at angles between 10°and 40°, where MWI is most sensitive to changes in tissue orientation, may exhibit paradoxical results if their orientation differs from the average orientation of the tract in the atlas. A difference in angle by 10°, for example, would result in a relative difference in MWF by approximately 5% to 7%, which is similar to the reduction in MWF in the normal appearing white matter in multiple sclerosis over five years (18). Similarly, if regions of interest are analyzed in a longitudinal study, participants should be scanned at the same head orientation. However, the fact that the dipole-dipole model provides a good fit to the experimental data suggests that an orientation correction based on that model and on an accompanying DTI scan may be feasible. Such correction would also be essential for the comparison of MWF between different fiber tracts.

In conclusion, our study revealed that the MWF depends on white matter fiber orientation. With increasing TR, the overall MWF decreased, while the orientation dependency persisted.

## Acknowledgments

This study was funded by the Austrian Science Fund (FWF) project number J 4038, by the National MS Society (RG 1507 05301), by the Natural Sciences and Engineering Research Council of Canada, Grant/Award Number 016-05371 and the Canadian Institutes of Health Research, Grant Number RN382474-418628. A.R. is supported by Canada Research Chairs 950-230363.

